# Improving PAH-chronically contaminated soil bioremediation using a combined strategy of bioaugmentation and surfactant-enhanced biostimulation

**DOI:** 10.1101/2025.09.09.675102

**Authors:** Esteban E. Nieto, Viviana A. Starevich, Laura Madueño, Irma S. Morelli, Coppotelli Bibiana, Festa Sabrina

## Abstract

Aged polycyclic aromatic hydrocarbon (PAH)-contaminated soil represents a challenge for the application of an effective bioremediation strategy, as the low PAH bioavailability limits microbial degradation. This study aimed to compare the effect of bioaugmentation (BA) and the coupled strategy of bioaugmentation and biostimulation (BA-SEB) on a chronically hydrocarbon-contaminated soil (IPK). For BA, we compared the performance of three PAH-degrading consortia with increasing diversity: SC AMBk, SC1, SC4. For BA-SEB, we coupled inoculation with these consortia with the addition of Triton X-100 surfactant, at sub-critical micelle concentration dose. In the BA microcosms, inoculation did not result in significant degradation of the determined PAH after 58 days of incubation. Among the BA-SEB microcosms, SC4 inoculated ones and the non-inoculated control showed significant degradation of fluoranthene, chrysene, and pyrene during the incubation period. Notably, BA-SEB microcosms inoculated with SC4 achieved superior pyrene degradation compared to the others. Both inoculation and the addition of surfactant impacted the community assembly, with the surfactant exerting the major effects. The surfactant addition stimulated several genera related to the degradation of PAH (*Novosphingobium, Blastomonas*). In addition, co-occurrence network analysis revealed that in SC4 inoculated microcosms most of these genera were positively correlated with the inoculated genera, particularly with *Paraburkholdeira* genus. The application of the combined strategy using the SC4 consortium in IPK soil successfully stimulated PAH biodegradation, demonstrating the importance of inoculant diversity and the identity of the consortium members under increased stress conditions (i.e from acute to aged contamination) for the strategy’s success.

## Introduction

Polycyclic aromatic hydrocarbons (PAH) are organic pollutants, with high toxicity, mutagenic and teratogenic properties, considered as priority control pollutants by the United States Environmental Protection Agency (EPA) (Patel et al. 2020). Soil is a major sink for PAH, where these compounds are gradually absorbed into the soil organic matter as the contamination time increases (Huang et al. 2024). Bioremediation strategies have been proposed as an eco-friendly and cost-effective approach to clean-up contaminated soils, since microbial processes are considered among the most significant in PAH removal (Miller et al. 2004; Patel et al. 2022). The bioavailability of PAH can be a main limitation for biodegradation, which can be affected by soil properties (Kuppusamy et al. 2017). Furthermore, aging processes, such as the absorption and sequestration of hydrocarbons within the soil matrix, pose an additional challenge for soil bioremediation. These processes contribute to increased stress by creating unevenly distributed pockets of concentrated pollution, degrading soil structure, and limiting the flow of oxygen and nutrients, all of which hinder microbial degradation (Haleyur et al., 2019; Curiel-Alegre, 2024).

Surfactant-enhanced biostimulation is a widely employed technology for aged PAH-contaminated soils (Kuppusamy et al. 2017; Lamichhane et al. 2017). Surfactants increase organic pollutants solubility by decreasing the interfacial surface tension between PAH and the soil/water interphase and by changing the partitioning effect of their micelles on the contamination ((Fernando Bautista et al. 2009; Gao et al. 2022). However, the high cost of surfactants and the potential for secondary pollution limit their application (Zhao et al., 2024; Wang et al., 2024). In addition, surfactants can also be preferentially utilized as carbon and energy sources for growth by PAH-degrading microorganisms, resulting in a decrease in metabolism of the target contaminant, thus inhibiting PAH biodegradation (Wolf et al. 2019). This is why, integrating diverse techniques on field-scale bioremediation has been proposed as a way to enhance the efficiency of remediating contaminated soil (Agnello et al. 2024; Wu et al. 2025). Several studies have shown that coupling bioaugmentation and surfactant addition can increase the pollutant removal rate in hydrocarbon-contaminated soils (Teng et al. 2022; Wu et al. 2025; Dai et al. 2025).

Bioaugmentation has shown positive results both in acute and aged PAH-contaminated soils (Brzeszcz et al. 2020; Wang et al. 2023; Nieto et al. 2025). Currently, the use of microbial consortia has gained popularity as potential inoculants. Positive interactions between their members can optimize the degradation processes (Massot et al. 2022), and increase stability and robustness, therefore, the probability of inoculum establishment (Stenuit and Agathos 2015; McCarty and Ledesma-Amaro 2019; Che and Men 2019). Recent studies have shown that the number of microbial species in an inoculant can be a critical parameter during bioremediation, enhancing its performance with higher diversity (Liu et al. 2023).

The outcome of the combined strategy (bioaugmentation and biostimulation) is still variable, and the effects on community structure are not well understood. Some reports have shown improvement in the degradation of hydrocarbons in contaminated soils mainly due to inoculation (Wolf et al. 2019), or due to surfactant application. The latter could act by stimulating the native degrading community (Ding et al. 2023) or even by causing inhibitory effects due to the selection of surfactant degrading bacteria. Coupling surfactant biostimulation with inoculation can alleviate the negative effects on the community structure caused by surfactant addition (Dai et al. 2025).

Many studies employ artificially spiked hydrocarbons, hindering understanding of real-world bioremediation for aged-contaminated soils. In previous works, the microbial community structure and functional potential of an aged-contaminated soil (IPK) was assessed through metagenomic analysis, demonstrating the presence of functional potential related to aromatic compound degradation (Festa et al. 2024). Allochthonous bioaugmentation with the PAH-degrading strain *Sphingobium AM* did not improve PAH removal in IPK soil (Festa et al 2016), whereas the application of the non-ionic surfactant Triton-X100 increased removal efficiency (Cecotti et al., 2018).

We have previously designed three PAH-degrading consortia with different number and composition of bacterial species (Macchi et al. 2021; Nieto et al. 2023), which achieved high PAH-degradation rates in freshly contaminated soil (Nieto et al. 2025). The purpose of this work is to explore the effects of these two bioremediation strategies (bioaugmentation and surfactant-enhanced biostimulation) on PAH degradation removal and on the native bacterial soil communities, to be used as a combined strategy. We hypothesized that this combined strategy will outperform bioaugmentation strategy in terms of PAH degradation and that it will exert higher impact on the bacterial soil community, and that the inoculant diversity is a major factor to be considered for the success of the combined strategy in PAH chronically contaminated soil. This study enhances our understanding of the coupled bioremediation strategies for enhancing contaminant removal in aged-contaminated soil.

## Material and Methods

### 2.1 Soil Samples

The aged contaminated soil (IPK) was collected from a petrochemical plant in Ensenada, Argentina (34° 53′ 19″ S 57° 55′ 38″W). This soil was previously treated by landfarming, with several applications of petrochemical sludge over 2 years. Soils were sampled ∼20 years after the closure of the treatment unit, showing a total PAH concentration of 573 ± 138 mg kg^-1^. More details on soil pollution history can be found in Festa et al (2024). Soil characterization was carried out using Bouyucus, Walkley–Black, Bray Kurtz and microkjeldahl methods for textural classification, organic carbon, available phosphorus and total nitrogen respectively. IPK soil was a loam soil with a pH of 7.71, 2.20% organic carbon, 3.78% organic matter, 0.20% total nitrogen, and 0.00083% available phosphorus.

### 2.2 Consortia preparation

The three defined consortia named SC AMBk, SC1 and SC4 were constructed with different combinations of the strains *Sphingobium* sp AM, *Paraburkholderia caledonica* Bk, *Pseudomonas* sp Bc-h and T, *Inquilinus limosus* Inq, and *Klebsiella aerogenes* B, all belonging to an enrichment culture previously obtained and characterised in the laboratory (Festa et al. 2013; Macchi et al. 2021). The populations responsible for the initial attack on phenanthrene are AM and Bk strains, while the other members (I, T, B and Bc-h strains) had the functional potential to degrade the pathway intermediates (Macchi et al. 2021). Each strain was grown independently in R3 broth at 28°C and 150 rpm overnight, then the cells were harvested by centrifugation at 6000 rpm and washed twice with sterile NaCl 0.85% (w/v). Cell number of each suspension was determined by measuring O.D at 580 nm. SC AMBk was composed of the strains AM and Bk in a 65:35 ratio, respectively; SC1 was composed of the strains AM, T, Bc-h, B and Inq; and SC4 of AM, Bk, T, Bc-h, B and Inq. Both SC1 and SC4 were built with equal proportions of each strain.

### 2.3 Microcosms set up

Microcosms of IPK soil were constructed with 150 g of sieved soil (2 mm mesh) in 350 ml glass containers. Two bioremediation strategies were carried out: bioaugmentation (BA) and a combined bioaugmentation and surfactant-enhanced biostimulation (BA-SEB). For BA, three treatments were performed by inoculating 5×10^7^ CFU/g dry soil of the 1) SC AMBk, 2) SC1 and 3) SC4 in IPK soil. For BA-SEB, sub-critical micelle concentration (sub-CMC) concentration of the nonionic surfactant octylphenol ethoxylate ether Triton X-100 (ultrapure, USB Corporation, USA) was supplemented (11 mg.g^-1^) as determined by (Cecotti et al. 2018) and the same inocula and concentration than in BA were used. Non-inoculated and non-biostimulated microcosms (IPK) and non-inoculated but Triton X-100 supplemented (IPK-T) were used as controls. The bioaugmentation strategy with the three consortia was also tested in IPK soil microcosms supplemented with phenanthrene (500 mg phenanthrene per kg dry soil), to assess if the physicochemical conditions of IPK soil were suitable for the degradation of a carbon source similar to contaminants, so only the PAH degradation analysis was performed. A non-inoculated but phenanthrene supplemented control (IPK-PHN) was also carried out.

Each microcosm was carried out in triplicates and incubated for 58 days at 24° C, and the final moisture was adjusted to 20% of humidity.

### 2.4 PAH quantification

PAH concentration was determined on 5 g of soil sampled from each microcosm at 0, 15, 30, and 58 days of incubation. Samples were lyophilized (L-3, REFICOR) and three consecutive extractions were carried out, using 15 ml of hexane:acetone 1:1 (v/v). The hydrocarbons were extracted in an ultrasonic bath (Testlab Ul-trasonic TB10TA) at 40 kHz, 400 W for 30 min. The mixture was centrifuged at 3000 rpm for 10 min and the supernatants were collected in brown glass flasks and evaporated. Then, samples were resuspended in 2 ml of hexane:acetone and filtered (0.45-μm nylon membrane). 5 μl of each sample was injected into a Perkin Elmer Clarus 500 GC-FID. The retention times of the different PAH were determined with standard solution and quantified using calibration curves through serial dilutions (Festa et al. 2024).

### 2.5 Sequencing Analysis

Total DNA for day 0 of control microcosms and for samples of all treatments after 30 days of incubation was extracted using E.Z.N.A. ® Soil DNA Kit repeating the lysis step twice to increase the extraction yield. Bacterial community composition was assessed using V4 hypervariable region of bacterial 16S rRNA gene using universal primer set 515F (GTGCCAGCMGCCGCGGTAA) and 806R (GGACTACHVGGGTWTCTAAT). Sample sequencing was performed using Illumina NovaSeq 6000 platform at Novogene Company. Paired end sequencing data was analyzed using QIIME 2 (v2022.2) (Bolyen et al. 2019). Raw data was filtered followed by denoising with DADA2 to obtain Amplicon Sequence variants (ASV). Taxonomy was assigned using SILVA v.138 trained database. Data analysis was performed using the *phyloseq* (v.1.42.0), *vegan* (v.2.6-4) and *Microviz* (v.) R packages. Prior to analysis, samples were rarefied to 106580 reads per sample (function *rarefy_even_depth*, with seed =1). To assess alpha diversity, Hill numbers were calculated after rarefaction, using the HillR package (version 0.5.2). Subsequently, beta diversity was analyzed through a Principal Coordinates Analysis (PCoA) using Bray-Curtis dissimilarity index at genus level.

The inoculated strains were monitored by ASV similarity as previously described in Nieto et al., (2025). Boxplots were used to study the relative abundance of the identified ASV for each inoculated strain across the studied treatments.

### 2.6 Co-ocurrence Networks

Co-occurrence networks were performed upon the main effect of inoculation at genus level. The networks were built using the Sparse Correlations for Compositional Data (SparCC) method using *microeco* R package (Liu et al. 2021). Only those correlations higher than 0.75 and a p-value lower than 0.05 were kept for the genera whose relative abundance was higher than 0.3%. Network attributes were calculated with *cal_network_attr()* function from the same package. Networks were visualized using Gephi v0.10.1

### 2.7 Statistical Analysis

All the tests were performed using rstatix (v.0.7.2) R package (Kassambara 2023). Statistical analysis of PAH concentration was performed through mixed ANOVA. Normality and homogeneity of variances assumptions were checked through Shapiro-Wilk and Levene tests, respectively. Pairwise comparisons were corrected using Benjamini-Hochman method. Differences in the diversity indices were assessed using Kruskal-Wallis test and Wilcoxon test for pairwise comparisons.

A distance-based variance partitioning analysis was performed to determine the amount of variance explained by the strategy and the treatment. Differential abundance analyses were carried out to identify the genera with significant changes between strategies and between inoculants in the combined strategy using the *ANCOM-BC* (v2.0.0) R package (Lin and Peddada 2020). A prevalence filter of 0.1, a significance of *p<*0.05 and multiple pairwise comparison considering the mdFDR, using holm as an adjusted method.

### 2.8 Data deposition

The raw data sequence for 16S rRNA gene amplicons has been deposited under NCBI accession number PRJNA1297662.

## Results

### 1. Influence of bioremediation strategies on PAH degradation

Degradation efficiency of the BA strategy and the combined BA-SEB strategy, both using three different consortia as inoculants, was evaluated by analysing the remaining concentration of PAH along incubation time (0, 15, 30 and 58 days) in each correspondent microcosms (Fig. 1). A non-inoculated and non-biostimulated microcosm was used as control (IPK).

**Figure 1.**
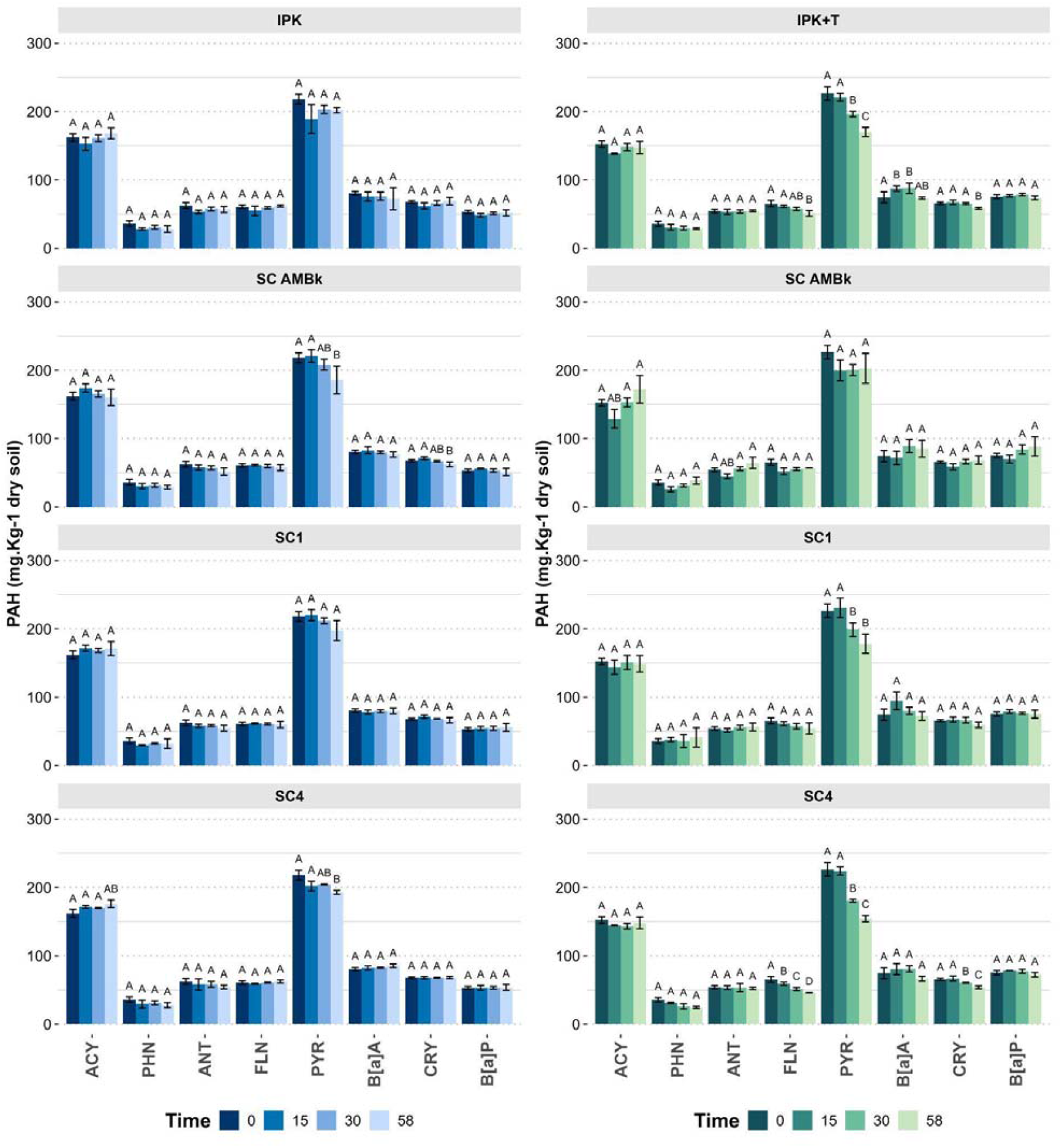
Quantification of PAH in IPK soil during bioaugmentation (BA, blue bars) and the combined strategy of bioaugmentation and surfactant-enhanced biostimulation (BA-SEB, green bars) at different incubation times during the 58 days period. The concentrations are expressed as the mean ± standard deviation. ACY: acenaphthylene, PHN: phenanthrene, ANT: anthracene, FLN: fluoranthene, PYR: pyrene, B[a]A: benzo[a]anthracene, CRY: chrysene, B[a]P: benzo[a]pyrene.

Among BA microcosms, inoculation with SC AMBk and SC4 consortia exhibited significant degradation of pyrene after 58 days (15% and 13% respectively), while no significant degradation of the other PAH was observed at the end of incubation period. No significant differences in PAH degradation were observed between SC AMBk and SC4 consortia compared to SC1 and control microcosms. To assess if the physicochemical conditions of IPK soil were suitable for the degradation, phenanthrene was supplemented to IPK soil and inoculated with the three consortia. The inoculated microcosms demonstrated significantly higher phenanthrene degradation (*p<*0.05) relative to the control IPK-PHN after 30 days of incubation (Fig S1). No other PAH degradation was observed during the incubation period.

Surfactant-enhanced biostimulation strategy (IPK-T) produced a significant degradation of fluoranthene (17%), pyrene (25%) and chrysene (12%) in relation to the non-inoculated and non-biostimulated microcosm (IPK) after 58 days of incubation. Comparable results were observed in microcosms inoculated with SC4, which also exhibited significant degradation of fluoranthene (30%), pyrene (32%), and chrysene (18%) (Fig. 1B). On the other hand, microcosms inoculated with SC1 only exhibited significant degradation of pyrene, while no degradation were observed in microcosms inoculated with SC AMBk.

Notably, among BA-SEB microcosms, after 30 days of incubation, only the combined strategy involving SC4 demonstrated superior pyrene degradation (20%, *p-adj <* 0.05) compared to the others. At the end of the incubation time, no significant differences were observed between IPK-T and SC4 microcosms, although a higher degradation trend was observed in the latter.

### 2. Impact of bioremediation strategies on bacterial community

To analyse the effect of the BA and the BA-SEB strategies on the microbial community, changes in alpha diversity were estimated through Hill numbers. No significant changes were observed between the diversity indexes of the inoculated microcosms of BA and BA-SEB (Fig S2). When comparing the effect of the strategy, no differences were observed in 0D and 2D (Fig. 2A and 2C). However, a decrease in 1D was observed in the combined strategy (*p*= 0.013) compared to BA (Figure 2B).

**Figure 2.**
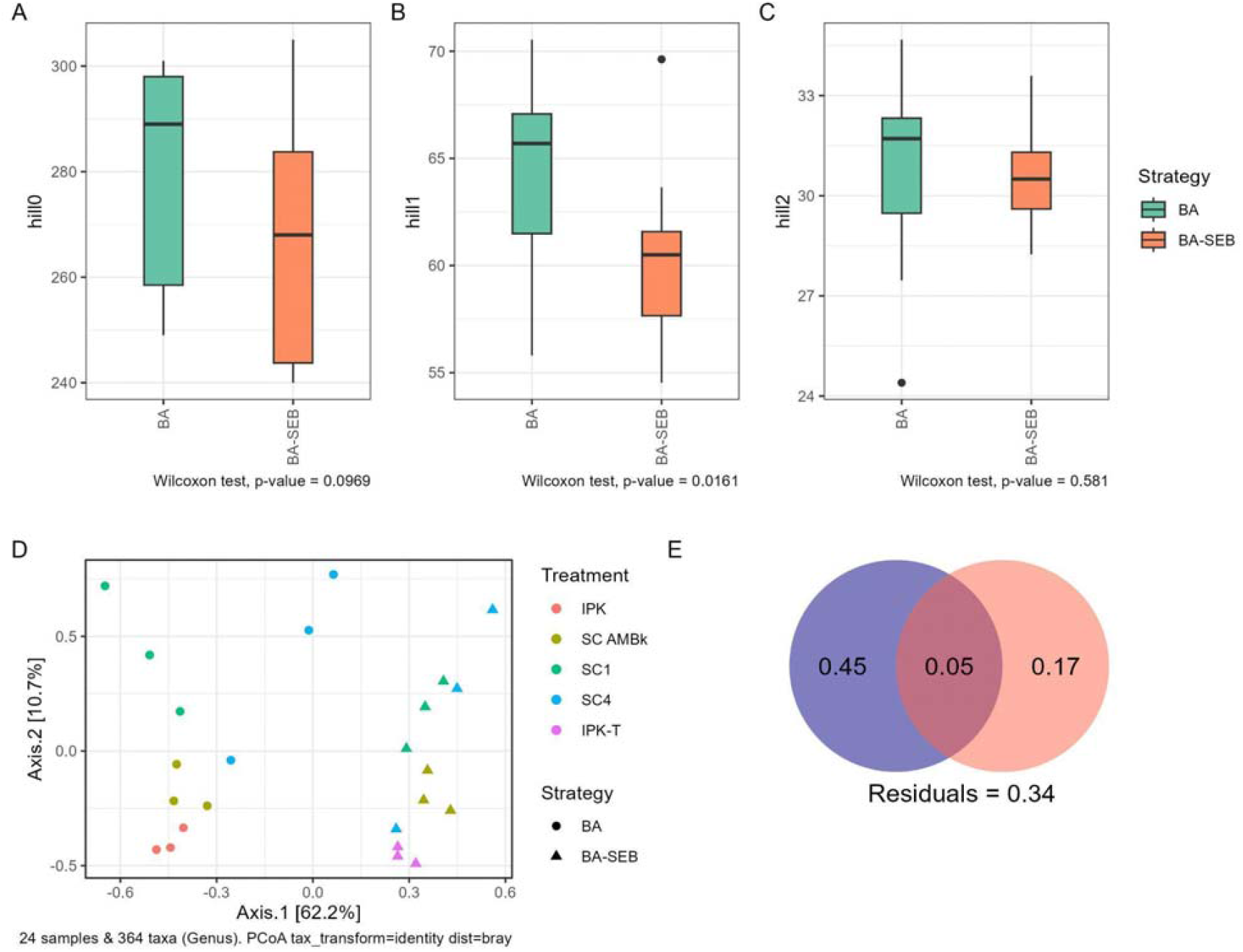
A) B) and C) alpha diversity analysis using Hill numbers comparing both strategies. D) beta-diversity analysis through a PCoA using Bray-Curtis dissimilarity index at genus level. E) distance-based variance partitioning analysis for the BA (pink) and BA-SEB (violet) strategies.

Regarding beta-diversity, the addition of Triton-X100 exerted an impact on community structure, as shown by the clustering analysis (Fig. S2) and PCoA (Fig. 2D). Analysis revealed two main clusters, where BA microcosms were grouped in the same cluster, except for BA-SC4 microcosms that were included together with BA-SEB microcosms. Both the addition of surfactant and the inoculation affected the community (dbRDA, *p<*0.01) which explained 66% of the variability (Fig. 2E). Most of this variability was explained by surfactant addition (45%), while the inoculation explained 17% of the variability.

### 3. Bacterial community shifts in the BA and BA-SEB microcosms

Community structure was analysed at genus level in BA and BA-SEB microcosms (Fig. 3). In both strategies, *Luteimonas, Immundisolibacter* and WN-HWB-116 were the predominant genera (Fig. 3A). Considering that the addition of Triton X100 was the main driver of variability in the community, ANCOM-BC was used to perform a differential abundance analysis of genera between both strategies (Fig. 3B). BA-SEB strategy mainly increased the abundance of 15 genera, most of them belonging to the Sphingomonadales order. Among the genera that showed higher change were *Blastomonas* and *Novosphingobium*, two members of Sphingomonadales order, and *Cavicella* and *Curpiavidus*, from Pseudomonadales and Burkholderiales orders respectively. Other members of Sphingomonadales order showed a similar trend, such as *Sphingobium* and a genus of Sphingomonadaceae family. Only two genera showed a significantly lower abundance in BA-SEB microcosms compared to BA, *Agromyces* and *S085*.

**Figure 3.**
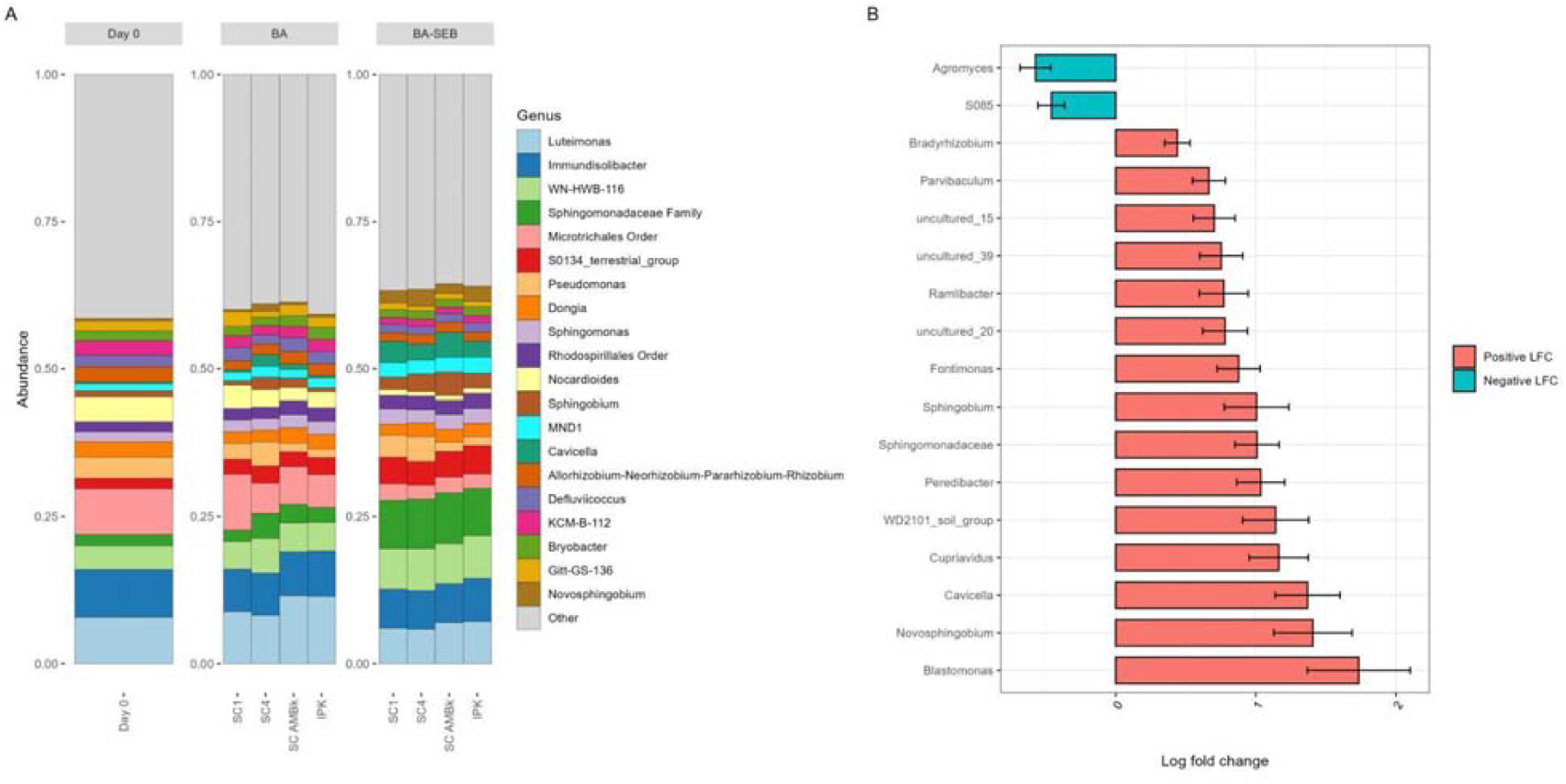
A) Relative abundance of the predominant genera at 30 days of incubation during bioaugmentation and the combined strategy of bioaugmentation and surfactant-enhanced biostimulation, expressed as the mean of the biological replicates. B) Log Fold-change (lfc) of the genera that showed significantly differential abundance between BA and BA-SEB strategies identified by ANCOM-BC.

### 4. Co-occurrence network analysis to study inoculation effect

To study the effect of inoculation on co-occurrence patterns of the bacterial community, a network analysis was performed at genus level. Table 1 shows the topological features of the networks. SC4 microcosms showed lower average degree, clustering coefficient and density, and longest average path length and heterogeneity. Also, SC4 microcosms showed the lower number of positive interactions (36.62%) compared to the other microcosms. Regarding the behaviour of the genera related to the inoculated strains, only the genus *Sphingobium* and *Pseudomonas* were present in the co-occurrence networks of all microcosms, while *Burkholderia-Caballeronia-Paraburkholderia* and *Enterobacteriaceae* family were present only in the networks of the microcosms that were inoculated with these strains (SC AMBk and SC4, and SC1 and SC4 respectively). A positive correlation was observed for *Sphingobium* and *Burkholderia-Paraburholderia-Caballeronia* when these two genera were present (Fig. 6A and B). The main differences were observed in the latter. The degree value of *Burkholderia-Paraburholderia-Caballeronia* in SC4 increased in comparison with the value of SC AMBk (30 and 14, respectively). In addition, positive correlations were observed between *Burkholderia-Paraburholderia-Caballeronia* and *Blastomonas, Novosphingobium, Sphingomonas, Bradyrhizobium* and *Xantobacteriaceae* family in SC4.

**Table 1.**
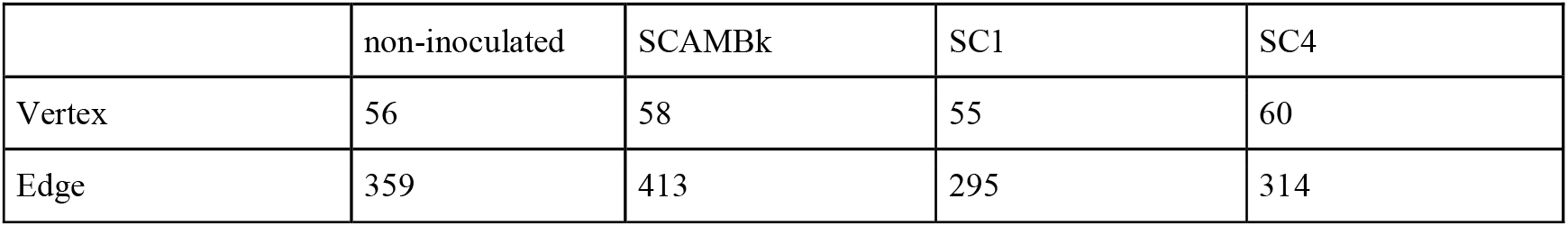
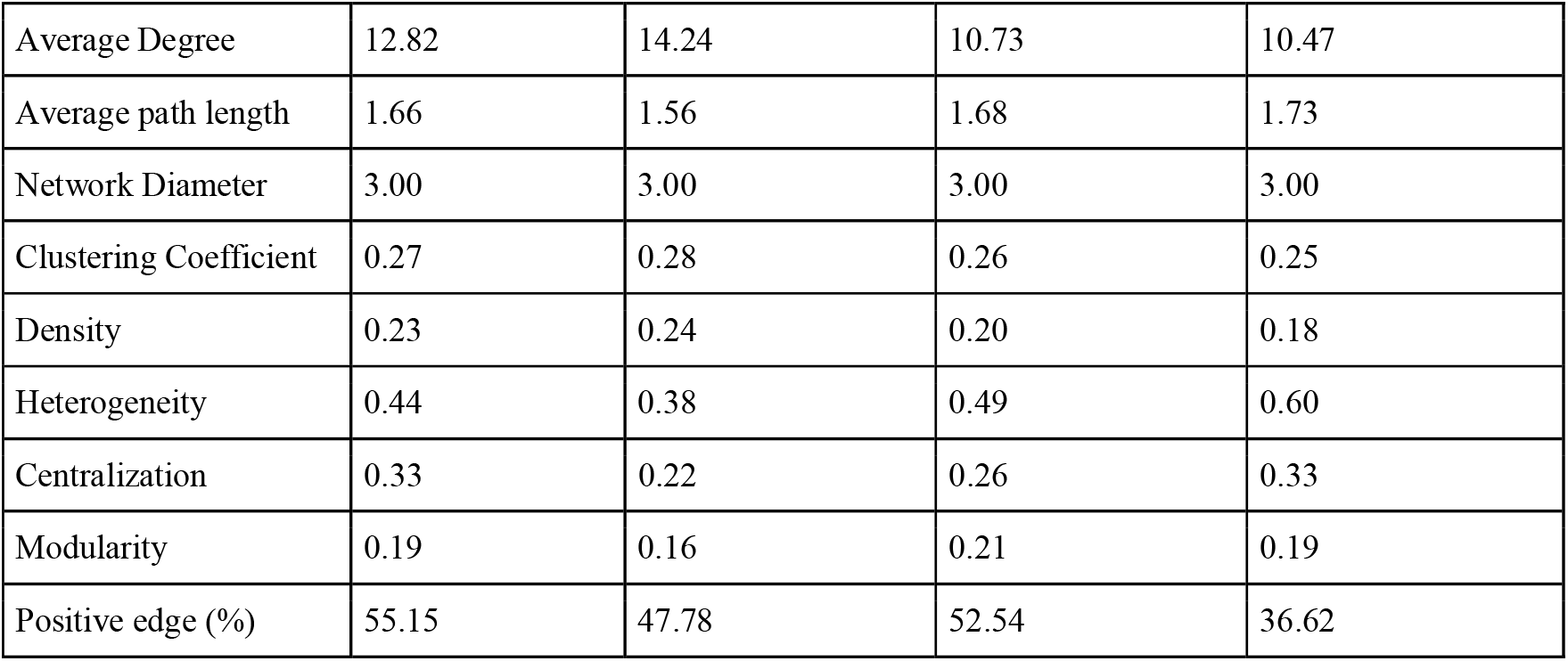
Topological properties of the co-occurrence networks.

### 5 Inoculant impact in BA-SEB microcosms

Given the differences in degradation efficiency observed among the inocula used in the BA-SEB microcosms, a differential abundance analysis using ANCOM-BC was performed at the genus level to assess the inoculant impact and compare the resulting changes on the microbial community composition (Fig. 5). SC AMBk inoculated microcosms showed a significant increase of the *Burkholderia-Caballeronia-Paraburkholderia*, and a decrease of the genera *Haliangium, UA11* and *uncultured 30*. Both SC1 and SC4 inoculated microcosms showed higher abundance of the inoculated genera *Inquilinus, Pseudomonas* and Enterobactereaceae family, while SC4 also showed a higher abundance of *Burkholderia-Caballeronia-Paraburkholderia*, compared with the SC1 microcosms.

The abundance of the ASVs identified for each inoculated strain was analyzed to track their fate in the bacterial community. Focusing on the BA-SEB strategy, microcosms inoculated with a consortia containing AM strain as a member (SC AMBk, SC1, and SC4) showed a higher abundance of the ASV identified for AM strain compared to IPK-T (Fig. S4). Microcosms inoculated with a consortia containing Bc-h and Inq strains as members (SC1 and SC4) exhibited a higher abundance of their corresponding ASVs than the other microcosms. However, the ASV identified for B showed greater abundance in SC1 compared to the other microcosms. For the Bk strain, its specific

ASV was most abundant in SC AMBk inoculated microcosms. However, among the other microcosms, it showed higher abundance in SC4 than SC1, but no significant difference was observed when compared to IPK-T (Fig. S4). Figure 6 compares the relative abundances of the ASVs identified for each strain between strategies. No significant differences were observed between the non-inoculated microcosms (i.e IPK and IPK-Triton). Notwithstanding, in the BA-SEB microcosms inoculated with SC AMBk higher abundance of all target ASVs were found when compared to BA strategy. Regarding the BA-SEB microcosms inoculated with SC1, a significantly higher (p< 0.05) relative abundance was found for AM and Bk related ASVs. On the contrary, Bk related ASV showed a significantly lower abundance in BA-SEB than in BA strategy.

## Discussion

Aged PAH-contaminated soil represents a challenge for the application of an effective bioremediation strategy, due to low bioavailability limits microbial degradation (Haleyur et al. 2019; Curiel-Alegre et al. 2024). Coupling surfactant enhanced biostimulation with bioaugmentation have shown promising results (Wolf et al. 2019; Teng et al. 2022; Wu et al. 2025; Dai et al. 2025). Here, we applied a combined strategy using the Triton-X100 surfactant and three defined consortia with different diversity. We compared the degradation efficiency and studied how each strategy affected the native microbial community. Our results showed that the addition of surfactant induced a change in the community structure (Fig. 2A, 2C, 3C), and that the combined strategy BA-SEB with the most diverse consortium (SC4), exhibited the highest degradation efficiency (Fig. 1) and increased the abundance of several PAH-degrading microbes from the native community (Fig. 3B).

Chronically contaminated soils represent an additional challenge to effective bioremediation, since limited resource availability makes it di□cult for the inoculants to establish and perform effective degradation (Haleyur et al 2019). Our previous results showed that inoculation of the strain AM or SC AMBk was not effective during PAH-degradation in IPK soil after 63 and 30 days of incubation, respectively (Festa 2016, Nieto 2024). Here we found that in the BA strategy, only SC AMBk and SC4 showed significant reduction of pyrene concentration after 58 days of incubation (Fig. 1A), with no significant differences compared to SC1 and control (IPK) microcosms.

The addition of surfactant increased PAH removal in the non-inoculated microcoms (IPK-T) (Fig. 1B), as previously reported in IPK soil by (Cecotti et al. 2018), which was mainly explained by an increase in the bioavailability of the sorbed fraction due to the addition of Triton X-100. Notably, during the combined BA-SEB strategy, a significant increase in the degradation of several PAH (pyrene, chrysene and fluoranthene) was demonstrated for the most diverse defined consortium (SC4) inoculated microcosms (Fig. 1B). This enhanced degradation of some PAH could be due to selective solubilization that can occur in systems containing diverse polycyclic aromatic hydrocarbons, potentially making different PAH more available for degradation (Bernardez and Ghoshal 2004; Masrat et al. 2013). Dai et al. (2025) reported similar results regarding PAH degradation in contaminated soil when applying biostimulation with Tween 80 and bioaugmentation with Aspergillus fumigatus, both individually and in combination, highlighting that the combined strategy significantly increased the PAH removal rate. Other studies have also shown synergistic effects coupling bioaugmentation and surfactant biostimulation (Curiel-Alegre et al. 2024; Wu et al. 2025). However, the outcome of the combined strategy with variable and non-additive effects was also reported (Wolf et al. 2019; Teng et al. 2022). Due to these inconsistent results, understanding the changes exerted in the native community by the individual and combined strategies is crucial. We observed that both the inoculation and the addition of the surfactant impacted the community structure, with the latter being the one that explained major variability in the community (Fig. 2 C and D). Analysis of the main effect of Triton-X100 addition revealed a decrease in the 1D Hill index, which could be a consequence of a change in taxa dominance caused by the surfactant (Wolf et al. 2019; Wu et al. 2025). In concordance with our results, (Teng et al. 2022) studied the active pyrene-degrading bacteria in pyrene contaminated soil microcosms using RNA-SIP during a bioaugmentation and a combined strategy with surfactant, and found that the number of active degraders increased in the combined strategy with no impact on the degradation efficiency. We also observed that the addition of Triton-X100 increased the abundance of several bacterial genera (Fig. 3B) known to be major PAH-degrading soil microorganisms, particularly some members of the Sphingomonadaceae family (including *Blastomonas, Novosphingobium, Sphingobium*) (Kertesz et al. 2019). Other well-known hydrocarbon degrading genera increased their abundance, such as *Cupriavidus, Fotnimonas, Cavicella, Parvibaculum* (Pérez-Pantoja et al. 2012; Fang et al. 2024; Chang et al. 2024). The observed increase in the abundance of the PAH-degrading microbes was already reported as one of the main mechanisms that contributes to enhanced removal of PAH (Wolf et al. 2019; Teng et al. 2021). The addition of surfactant can also induce changes in the abundance of microbes that consume the added surfactant using it as a preferential carbon source, inhibiting the degradation of the target pollutants (Ghosh and Mukherji 2016; Cecotti et al. 2018; Wolf et al. 2019). These negative effects of surfactant addition can be alleviated by combining its application with bioaugmentation (Dai et al. 2025).

Inoculation also had a significant impact on the community structure (Fig. 2 C and D). In BA strategy, the community structure present in the microcosms inoculated with SC4 was more similar to the one of the BA-SEB systems (Fig. 2B and S2) and, exhibited the highest PAH-degradation rate in BA-SEB strategy compared to the other inoculated microcosms. The inoculation with the defined consortia, modified the co-occurrence patterns of the soil community (Fig. 6). In particular, in SC4 inoculated microcosms the number of negative edges was increased (Table 1, Fig. 6D). Competitive interactions introduce negative-feedback loops that have a stabilizing effect, increasing the resilience of the community (Coyte et al. 2015). Positive correlations were observed between the genus *Sphingobium* and *Burkholderia-Paraburkholderia-Caballeronia* in SCAMBk and SC4 inoculated microcosms, as previously reported for PAH acute contaminated soil (Nieto et al. 2025). Notably, the degree of *Burkholderia-Paraburkholderia-Caballeronia* in SC4 inoculated microcosms was higher than in the other microcosms and positive correlations were observed between this genus and *Blastomonas, Novosphingobium, Sphingomonas, Bradyrhizobium*, among others. Most of these genera were also stimulated by the addition of the surfactant (Fig. 3B). Previous reports have shown that inoculation can increase the PAH removal by changing the co-occurrence pattern and stimulating the PAH-degrading community, even when the inoculum does not actively participate in the degradation (Li et al 2018, Teng et al 2022). Thus the increment of the degradation efficiency in BA-SEB inoculated with SC4 could be explained as a synergistic effect of the inoculant and the surfactant in the selection of PAH-degrading bacteria from the native community.

The use of allochthonous microbes as inoculants offers the advantage of reducing time and cost, and it has been proven successful for PAH removal (Brzeszcz et al. 2020; Nieto et al. 2025). However, one of the main challenges is to ensure the survival of the introduced microbes, which should face new abiotic and biotic barriers that can limit the establishment (Albright et al. 2022). Our previous results showed that bacterial predators respond quickly to inoculation, reducing inoculant survival and affecting PAH degradation when the growth rate is lower than the predation rate (Nieto et al. 2024). The addition of surfactant led to an increased abundance of the inoculated genera (Fig. 5), which can be attributed to the increase in PAH bioavailability. This improved resource access directly contributed to the inoculum’s enhanced survival. However, contrary to our previous results in acutely-contaminated soil (Nieto et al. 2025), where the degradation efficiency did not depend on consortium diversity, as discussed above we observed a significant difference regarding the degradation performance of the different consortia. This study showed that the most diverse consortium exhibited the highest degradation performance in aged-contaminated soil. Although these two results seem contradictory, they are consistent with the results reported by (Fetzer et al. 2015). These authors found that benzoate degradation by artificial communities varies in environments with increasing abiotic stresses. While in low stress conditions these communities performed well even with low richness, no such performance was observed in more stressed conditions, where the addition of new species led to an increase in community function (Fetzer et al. 2015). Additionally, these authors showed that the function was dependent on species identity and that the interactions among the members also changed with greater stress, increasing the interdependence among the species. In agreement with these findings, our results showed that both the species identity and the diversity of the consortia play a role in the degradation efficiency, since SC4 not only included the highest diversity but also included both PAH-degrading strains (AM and Bk). In SC4 inoculated microcosms, the presence of Bk was key to enhance the degradation, since it played a role modulating the co-occurrence pattern (Fig. 4D). However, inoculating only the PAH-degrading bacteria was not enough, since SC AMBk failed to enhance PAH-degradation and neither SC1 (harboring a single active degrader) improved PAH degradation. As in Fetzer et al (2015), when the stress increases (i.e. from acute PAH contamination to aged-PAH contamination) the interdependence among the consortium members also increases.

**Figure 4.**
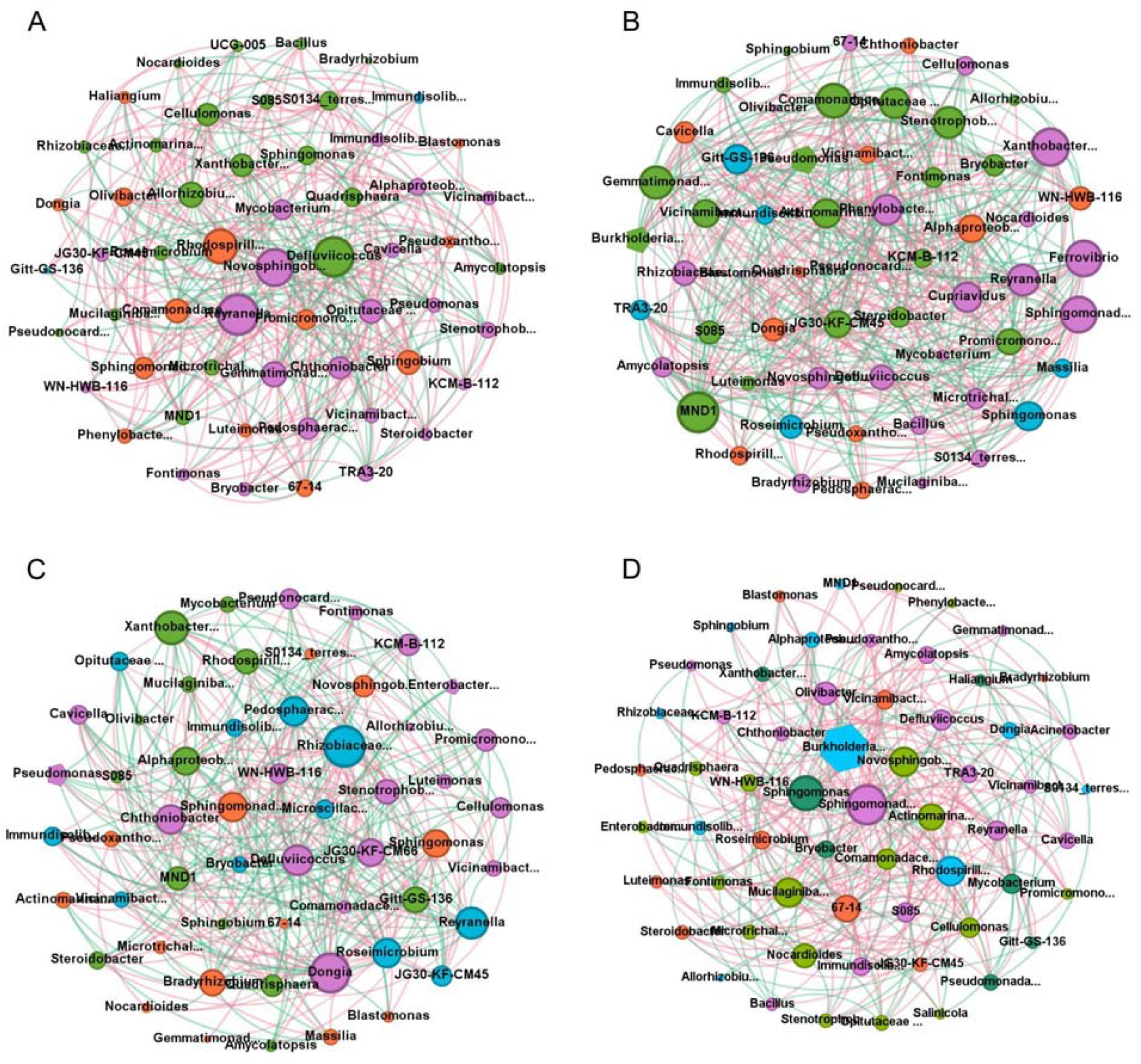
Co-occurrence networks at genus level to assess the main effects of inoculation, using SparCC method for IPK soil microcoms A) Control, B) SC AMBk, C) SC1 and D) SC4. Only significant correlation (p<0.05) higher than 0.75 are shown. The colors represent the different identified modules and the size of the nodes represent the number of direct connections that node has to other nodes in the network (degree value). Red and green edges represent negative and positive correlations, respectively. Inoculated genera nodes are represented as pentagons.

**Figure 5.**
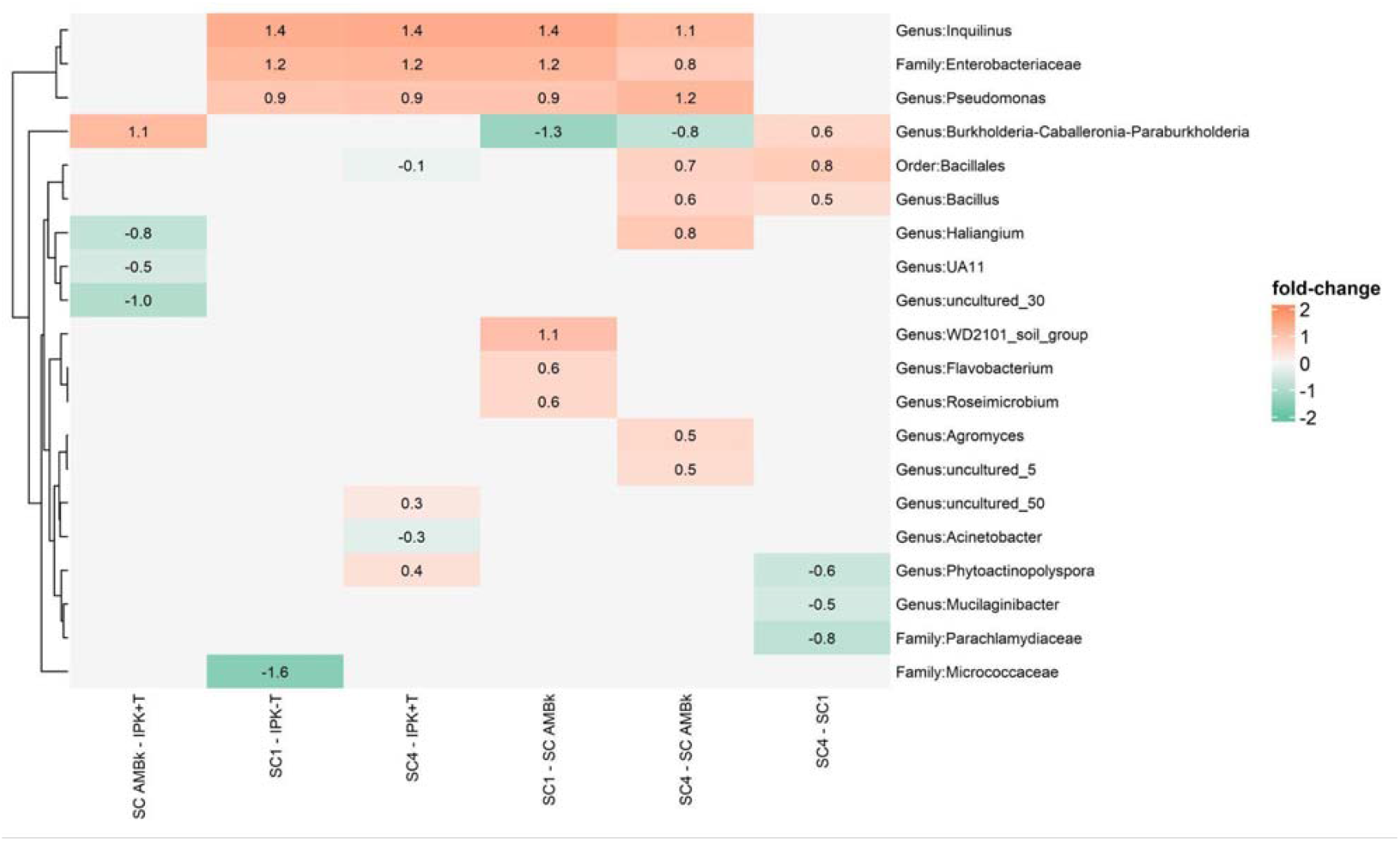
Fold-change of genera with significant differential abundance, as identified by ANCOMBC, between the different inoculated microcosms of the BA-SEB strategy.

**Figure 6.**
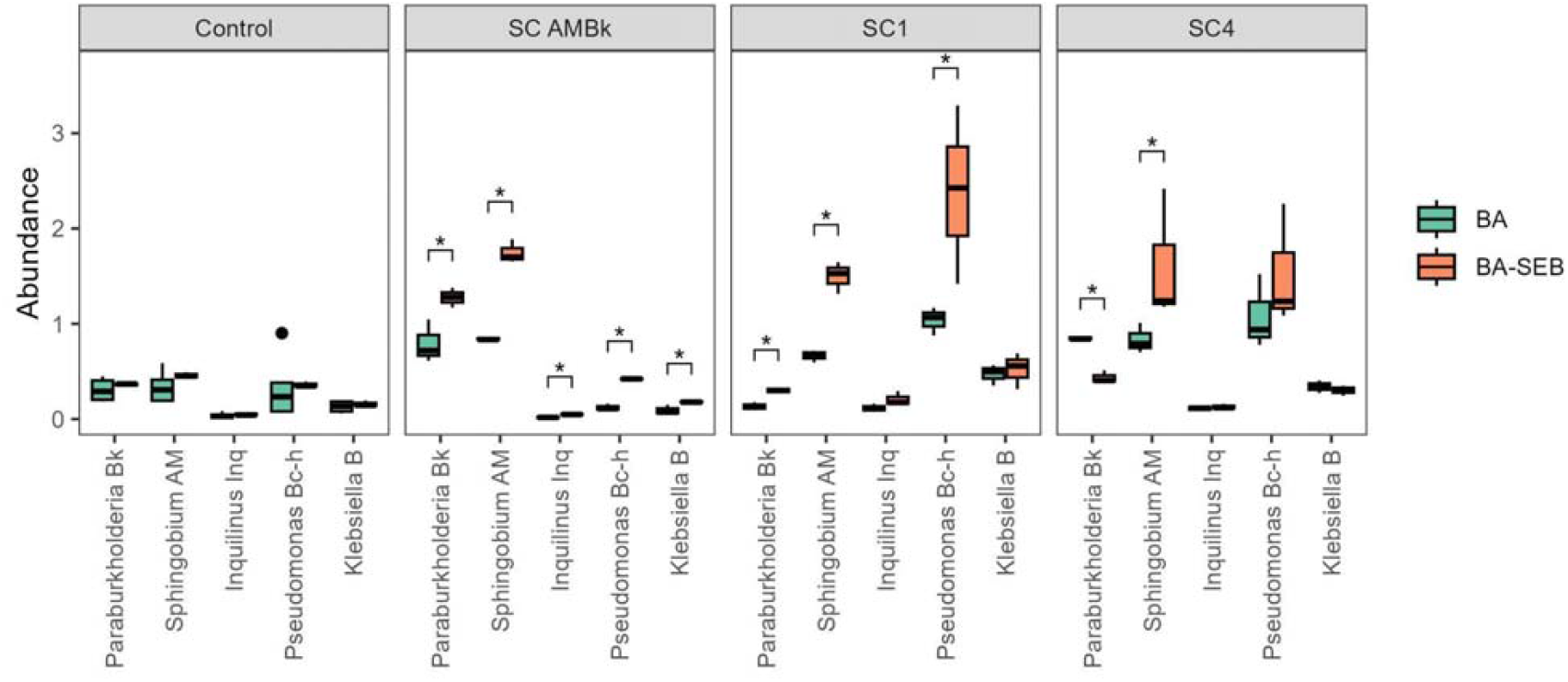
Relative abundance of the ASVs identified for each inoculated strains for both BA and BA-SEB strategies.

In our work the combined strategy exerted a synergistic effect, increasing the degradation of several PAH. However, residual PAH concentration remained high at the end of the incubation period. To address these limitations, some authors have improved the efficiency of the combined strategy by re-inoculation (Curiel-Alegre et al. 2024; Wu et al. 2025) or by the addition of the inoculant through carrier materials (biochar, montmorillonite), increasing the half life of the inoculants and shielding microbes from environmental stresses (Wang et al. 2023; Curiel-Alegre et al. 2024). These parameters are key in to maximise inoculum survival and thus enhance the degradation efficiency in combined strategies.

## Conclusion

We demonstrated that the combined strategy of bioaugmentation using the SC4 bacterial consortium along with surfactant-enhanced biostimulation in IPK soil led to a significant increase in PAH biodegradation. The improved degradation efficiency observed in this combined strategy suggests a synergistic effect between the inoculant and the surfactant, which positively impacted the assembly of the bacterial community. This process promoted the selection of native PAH-degrading bacteria. Furthermore, our findings highlighted that the diversity of the inoculant and the specific identities of the consortium members are crucial for successful bioremediation in aged PAH-contaminated soils under stressful conditions.

## Acknowledgments

This work was financially supported by Consejo Nacional de Investigaciones Científicas y Técnicas (CONICET), Agencia Nacional de Promoción Científica y Tecnológica (PICT 2019-1805; PICT 2019-2540).

## Author contribution

**Esteban E. Nieto**: Conceptualization, Investigation, Formal analysis, Visualization, Writing - Original Draft, Writing - Review & Editing; **Viviana A. Starevich**: Formal analysis, Writing - Review & Editing; **Laura Madueño**: Visualization, Writing - Review & Editing; **Irma S. Morelli**: Funding acquisition, Project administration; **Bibiana M. Coppotell**i: Conceptualization, Funding acquisition, Project administration, Writing - Review & Editing; **Sabrina Festa**: Conceptualization, Supervision, Funding acquisition, Writing – Original Draft, Writing - Review & Editing.

## Notes

### Competing Interest Statement

The authors have declared no competing interest.

