## Supplemental material for "Improving PAH-chronically contaminated soil bioremediation using a combined strategy of bioaugmentation and surfactant-enhanced biostimulation"

**Supplementary material**

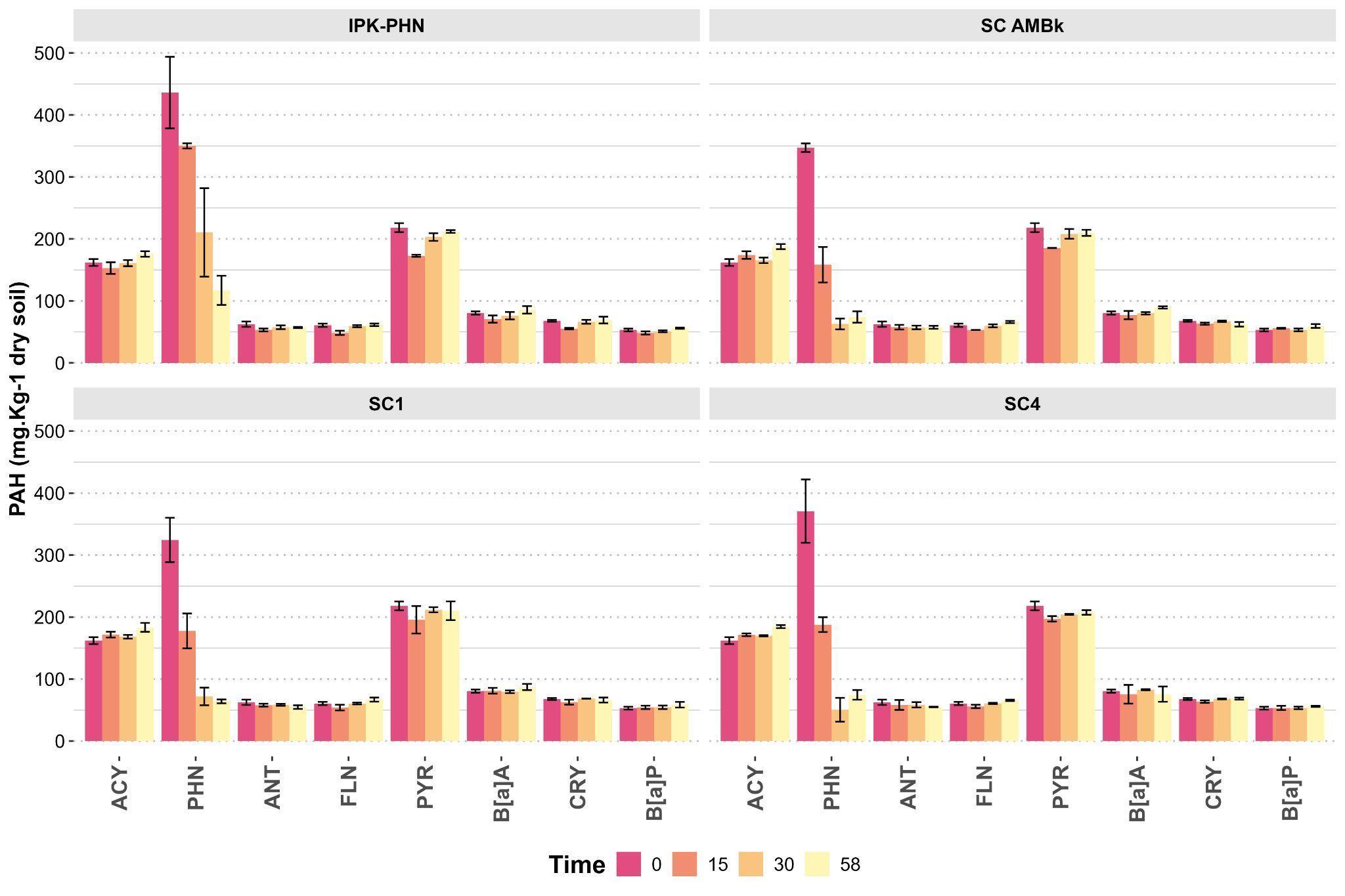

**Figure S1**: Quantification of PAH detected in IPK soil after the addition of PHN as a carbon source, during bioaugmentation strategy at different incubation times during the 58 days period. The concentrations are expressed as the mean ± standard deviation. ACY: acenaphthylene, PHN: phenanthrene, ANT: anthracene, FLN: fluoranthene, PYR: pyrene, B[a]A: benzo[a]anthracene, CRY: chrysene, B[a]P: benzo[a]pyrene.

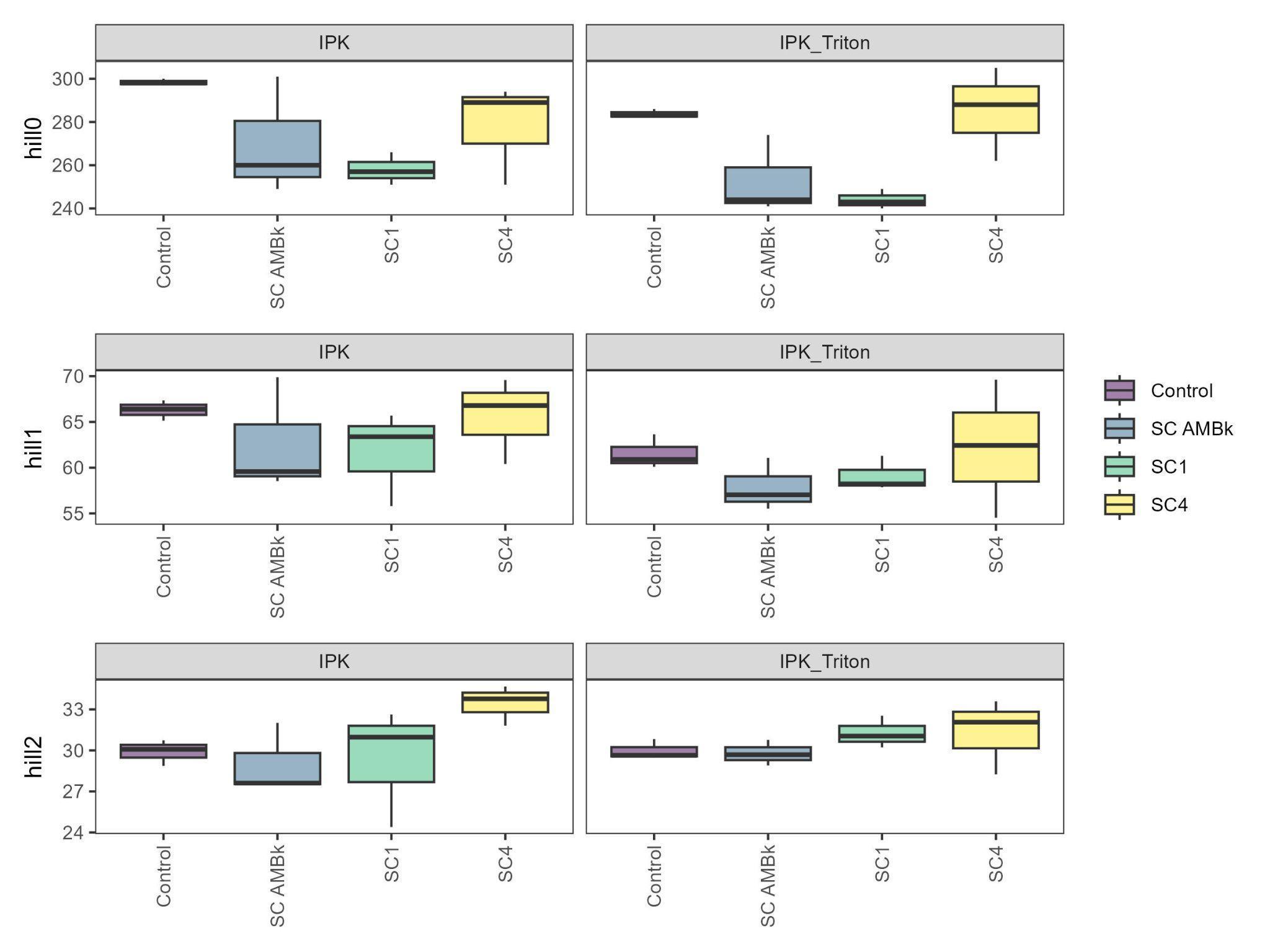

**Figure S2:** alpha diversity of BA and the BA-SEB microcosms estimated through Hill numbers

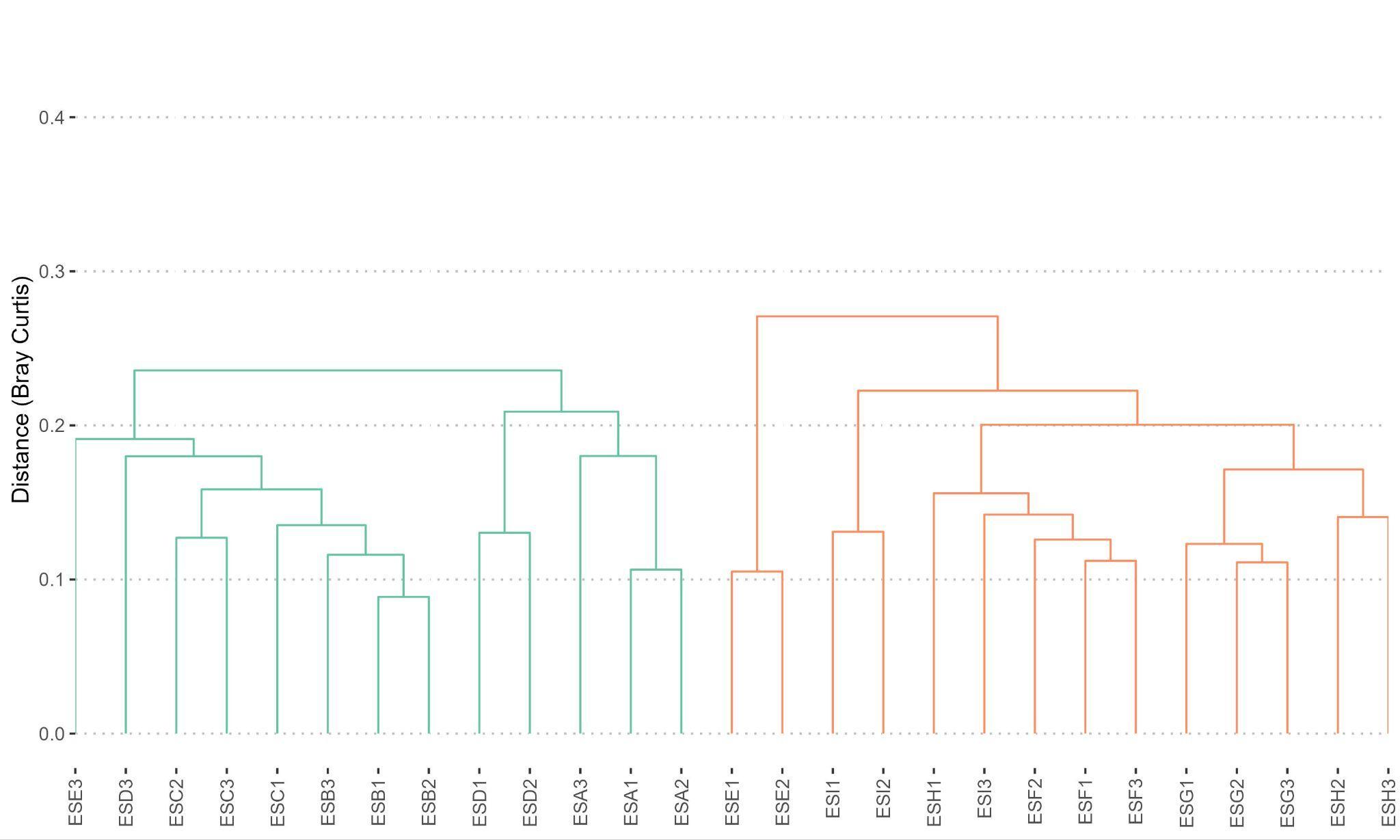

**Figure S3:** Clustering analysis of BA and the BA-SEB samples using Bray-Curtis distance and Ward methods. BA: IPK control (ESA1, ESA2, ESA3, ESB1, ESB2, ESB3); CS AMBK (ESC1, ESC2, ESC3); CS1 (ESD1, ESD2, ESD3), CS4 (ESE1, ESE2, ESE3). BA-SEB: IPK-T (ESF1, ESF2, ESF3); CS AMBK (ESG1, ESG2, ESG3); CS1 (ESH1, ES2, ESH3); CS4 (ESI1, ESI2, ESI3).

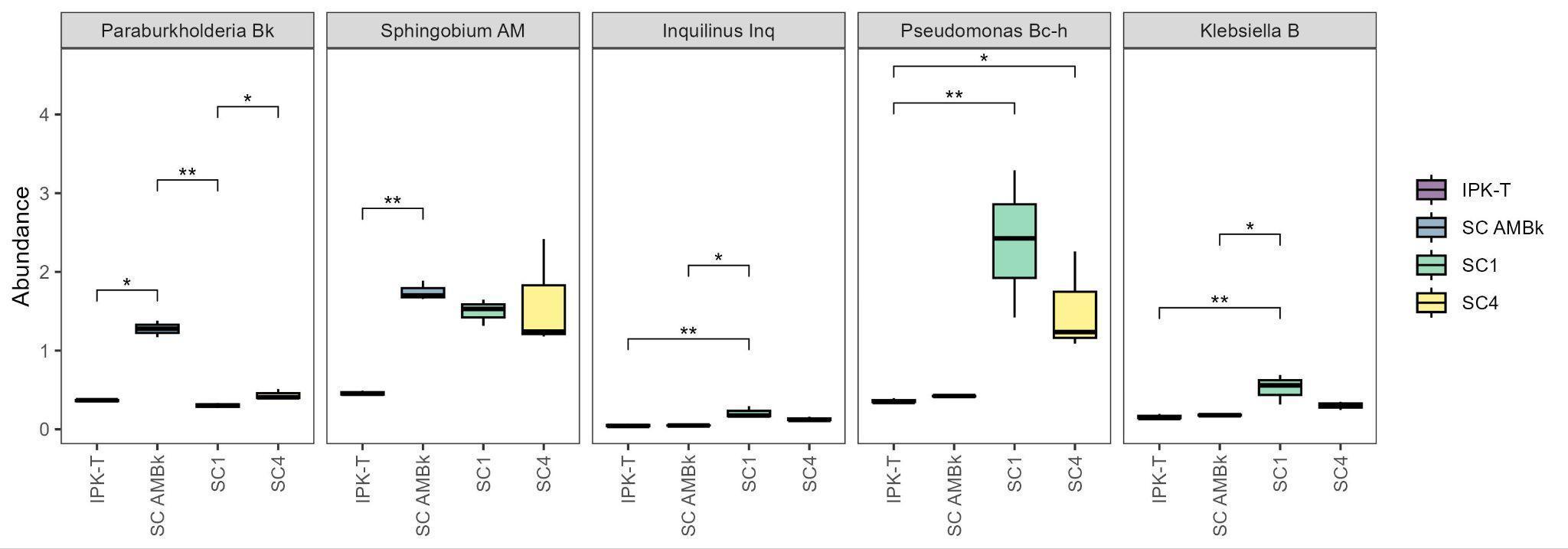

**Figure S4:** Relative abundance of the ASVs identified for each inoculated strains in BA-SEB strategy
